# Differential effects of opioid receptor modulators on motivational and stress-coping behaviors in the back-translational rat IFN-α depression model

**DOI:** 10.1101/769349

**Authors:** Charlotte K. Callaghan, Jennifer Rouine, Md Nurul Islam, David J. Eyerman, Karen L. Smith, Laura Blumberg, Connie Sanchez, Shane M. O’Mara

## Abstract

**Rationale:** Many patients respond inadequately to antidepressant drug treatment; the search for alternate pharmacological treatment mechanisms is ongoing. Until the 1950’s, opium was sometimes used to treat depression, but eventually abandoned due to addiction risk. Recent insights into opioid biology have sparked a renewed interest in the potential antidepressant properties of opioids.

**Objective:** We studied how mu (MOR), kappa (KOR) and delta (DOR) opioid receptor ligands affect the dysregulation of motivated behavior (progressive ratio responding; PR), stress-coping behavior (forced swim test; FST) and hippocampal neurogenesis in rats, all induced by the back-translational interferon-alpha (IFN-α)-induced depression model.

**Methods:** Male Wistar rats (3-months old, 8/group) were treated with recombinant human IFN-α (170,000 IU/kg, 3 times/week) or saline. Ligands of the MOR, KOR and DOR receptors were administered as follows: a single subcutaneous dose, 30min before PR and 1h before FST, of the MOR agonist morphine (full agonist; 5mg/kg), the partial agonist RDC 2944 (0.1mg/kg) and the antagonist, cyprodime (10mg/kg); of the KOR agonist, U50 488 (5mg/kg), the antagonist, DIPPA (10mg/kg); and the DOR agonist, SNC 80 (20mg/kg) and antagonist naltrindole (10mg/kg). After 4 days of treatment with the mitotic BrdU marker, hippocampi were harvested and analysed for neurogenesis. Fluoxetine (10 mg/kg/day for 4 weeks, orally) served as control for assay sensitivity in the FST.

**Results:** The KOR antagonist, DIPPA, the DOR agonist SNC 80 and fluoxetine reversed the IFN-α-induced immobility increase in the FST. The MOR agonist, morphine, the KOR antagonist DIPPA, and the KOR agonist U50 488 reduced the IFN-α-induced increase in the breakpoint in the PR. The DOR agonist SNC 80 recovered the IFN-α-induced decrease in BrdU+ hippocampal cells.

**Conclusion:** Opioid receptors mediate different aspects of the IFN-α-induced dysregulation of motivational and stress-coping behaviors and hippocampal neurogenesis in a back-translational model of depression. KORs and DORs appear to play more prominent roles in torpor–inertia-type behaviors, whereas DORs appear more involved in the regulation of neurogenesis.

## 1. Introduction

The World Health Organization reports more than 300 million people are living with Major Depressive Disorder (MDD). Standard antidepressant therapies, including selective serotonin reuptake inhibitors (SSRIs), require treatment for several weeks to provide efficacy; about two-thirds of patients are poor or non-responders to drug treatment [1], [2]. Therefore, there is a major unmet need for novel antidepressant therapies. Preclinical research to identify novel antidepressant mechanisms is challenging since MDD is a heterogeneous multifactorial disorder that presents itself clinically with variable symptomatology between patients [2]. For these reasons, preclinical investigations related to MDD must be limited to mimic particular symptoms rather than the clinical syndrome.

Opium was until mid-last century commonly used for treating depression, but abandoned following the introduction of the monoaminergic antidepressants, due to the inherent risk of addiction [3]. However, advances in understanding opioid neurobiology have sparked renewed interest in opioids as antidepressants. The endogenous opioid peptides (endorphins, enkephalins and dynorphins) and their corresponding G-protein coupled receptors (GPCR) (mu, kappa and delta opioid receptors (MOR, KOR and DOR, respectively)) are integral to regulation of motivation and stress responses [4]; their expression pattern heavily overlaps with brain regions implicated in motivational and stress-coping behaviors [5–7]. Opioid receptors are extensively expressed in brain regions implicated in MDD, including the hippocampus (HPC), nucleus accumbens (NAc), medial prefrontal cortex (mPFC), amygdala, claustrum, thalamus, hypothalamus, ventral tegmental area (VTA) and the dorsal raphe nucleus (DRN) [5–7]. MORs have been implicated in motivational and anxiety-like behaviors in the forced swim test (FST), open field and elevated plus maze. [8, 9]. Clinical studies suggest the therapeutic potential of opioid receptor modulators for MDD [3, 10–12]. The partial MOR agonist, buprenorphine, alone or in combination with the MOR antagonist, samidorphan, improved HAMD scores in patients with MDD or improved emotional responses in patients with a range of mood symptomatology. Further, administration of synthetic β-endorphins improved behavioral scores in depressed patients in just two to four hr following treatment.

The present work focuses on opioid receptor modulators in dysregulated motivational and stress coping behavior [13, 14], using the clinically-relevant, back-translational, rat interferon-alpha (IFN-α) depression model [15–19]. We employed a progressive ratio (PR) task reinforced with a natural reward (sucrose pellets) to examine motivated behavior, and the FST to assess stress-coping behavior. In addition to the behavioral measures, effects on IFN-α-induced reduction of hippocampal neurogenesis were assessed using flow cytometry analysis. The following opioid receptor modulators were investigated: morphine, RDC 2944 ((4bR,6S,8aS,9R)-11-(cyclobutylmethyl)-4,6-dihydroxy-8a-methoxy-6,7,8,8a,9,10-hexahydro-5H-9,4b-(epiminoethano)phenanthrene-3-carboxamide hydrochloride), cyprodime (MOR full agonist, partial agonist and antagonist, respectively), SNC 80, naltrindole (DOR agonist and antagonist, respectively), U50 488 and DIPPA (KOR agonist and antagonist, respectively). RDC 2944 is a novel selective MOR agonist with low intrinsic activity; the basic pharmacology profile is summarized in section 2.1. The compounds were chosen based on their specific pharmacology, high affinity for their associated receptors and limited interactions with other receptor systems [20–25].

## 2. Materials and Methods

### 2.1. Pharmacological profile of RDC 2944

RDC 2944 was provided by Alkermes, Inc. Radioligand receptor binding assays were performed to determine the affinity for each of the opioid receptor types (MOR, KOR and DOR). The binding assays were performed using the method described in detail in [26]. The Ki values for MOR Ki =0.42 nM; KOR Ki = 9.1 nM and DOR Ki = 84 nM were calculated according to Cheng and Prusoff [27].

The [35S]GTPγS binding assay was used to determine the agonist and antagonist properties of RDC 2944. The [35S]GTPγS binding assay was performed as previously described in detail in Bidlack and colleagues [26]. The concentration needed to achieve 50% of maximal MOR stimulation by RDC 2944 (EC50) = 8.7 nM, and the maximal MOR stimulation over basal conditions (E_max_) = 16 %. KOR EC50 = 46 nM; KOR E_max_ = 37% were also measured. To determine the MOR antagonist properties of RDC 2944, membranes were incubated with varying concentrations of DAMGO. The concentration needed to achieve 50% of maximal MOR inhibition by RDC 2944 (IC_50_) = 15 nM and the maximal MOR inhibition over basal conditions (I_max_) = 87 %. Together, these data indicate that RDC 2944 is a selective low intrinsic activity agonist for the MOR.

Dose selection for *in vivo* testing was determined following preliminary pharmacokinetic and receptor occupancy analysis. Using *ex vivo* autoradiography (method in supplement), RDC 2944 occupancy at MOR was determined to be zero at 0.03 mg/kg (-0.2 ± 12%) and achieved a maximum of 67 ± 3.8% receptor occupancy at 1 mg/kg 1hr post sc administration (Figure S1). RDC 2944 occupancy at KOR was effectively zero (%) at all doses tested. RDC 2944’s ED50 and hillslope were not calculated due to the restricted range of occupancy values achieved at the doses used here. Total brain exposure was determined using LC/MS/MS (Method in supplement). Acute sc administration of RDC 2944 induced dose-related increases in total brain exposure, ranging from 1.4 ± 0.4 nmol/kg at 0.01 mg/kg to 110 ± 23 nmol/kg at 1 mg/kg 1hr post dose (Figure S2).

### 2.2 Behavioral studies

#### 2.2.1. Animals

Male Han Wistar rats (421 ± 5g: weights on arrival), sourced from Envigo UK, were used in these experiments. The animals were pair-housed in a controlled environment (temperature: 20–22°C, 12/12 hr light/dark cycle), with water and food *ad libitum* (unless otherwise stated), and maintained under veterinary supervision. Animal weights were monitored throughout the experiment; no changes between groups were observed (data not shown). All experiments were performed under license from the Health Products Regulatory Authority (HPRA) of Ireland in accordance with EU regulations and with local ethical approval (the University of Dublin, Trinity College Dublin), license number AE19136/P051.

#### 2.2.2. Dosing regimens

Dosing regimens are summarized in figure 2.

**Figure 1.**
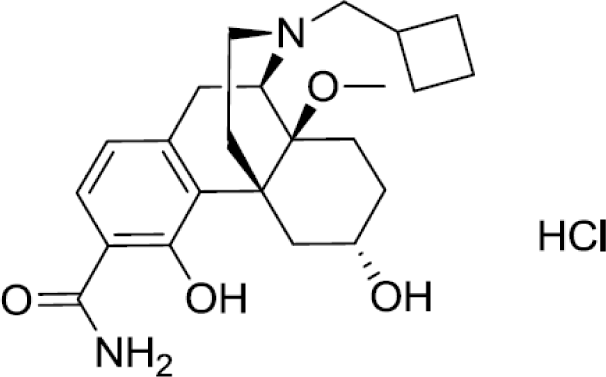
Chemical Structure of RDC 2944.

**Figure 2.**
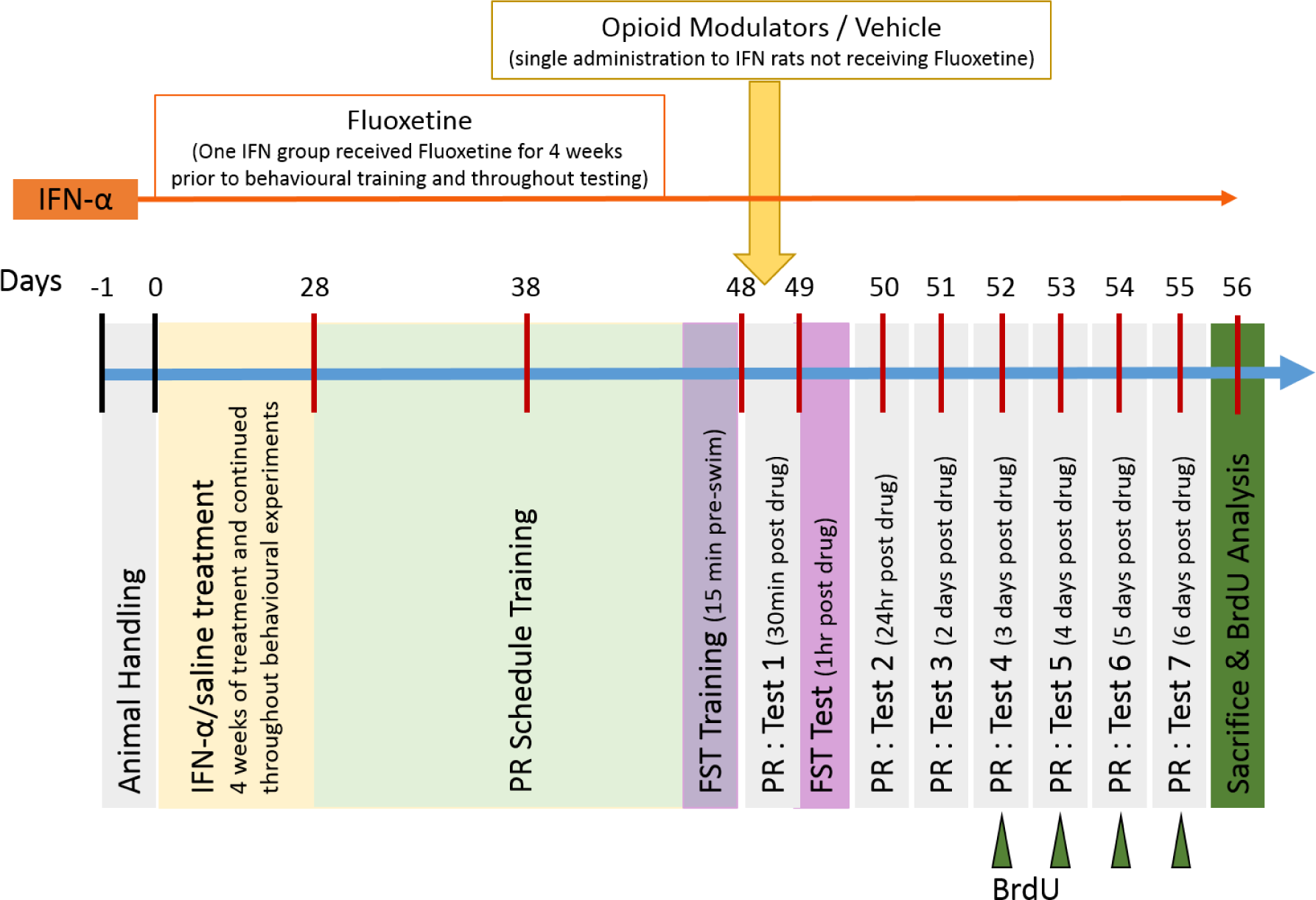
Experimental design: IFN-α or saline (control) was administered 3 times per week for 4 weeks before starting the behavioral experiments and continued treatment for the duration of the experiments. Animals treated with fluoxetine (10 mg/kg suspended in maple syrup po) were treated daily for 4 weeks prior to commencing behavioral experiments and continued treatment for duration of the experiments. Cyprodime (10mg/kg), morphine (5mg/kg), RDC 2944 (0.1mg/kg), DIPPA (10mg/kg), U50 488 (5mg/kg), naltrindole (10mg/kg) or SNC 80 (20mg/kg)) were administered 30min prior to test in the PR schedule and 1hr prior to test in the FST on day 49. Animals were repeatedly tested in the PR schedule for 6 days following drug treatment. To investigate neurogenesis, animals received injections of BrdU (100mg/kg) on days 52-55 as indicated by solid green arrows. On day 56 animals were sacrificed and tissue was harvested for BrdU analysis by flow cytometry.

IFN-α treatment was administered as previously published [15, 16, 18]. Briefly, the rats were treated with either 170,000 IU/kg IntronA (human recombinant interferon-alpha 2b, Merck Sharp & Dohme Limited), or saline (0.9% NaCl) three times per week for four weeks before starting behavioral testing, and throughout the remaining duration of the study. Injection solutions were prepared in saline and administered at a volume of 1 ml/kg subcutaneously (sc). IFN-α / saline treatments were administered approximately 24hr before behavioral testing.

Opioid Receptor Modulators

MOR agonists: morphine (5 mg/kg; Martindale Pharmaceuticals), and RDC 2944 hydrochloride (HCl) (0.1 mg/kg; Alkermes, see pharmacology in section 2.1), both dissolved in saline.

MOR antagonist: cyprodime HCl (10 mg/kg; Tocris) dissolved in 10% dimethyl sulfoxide (DMSO).

KOR agonist: (±)-U50 488 HCl (10 mg/kg; Tocris) dissolved in saline.

KOR antagonist: 2-(3,4-Dichlorophenyl)-N-methyl-N-[(1S)-1-(3-isothiocyanatophenyl)-2-(1-pyrrolidinyl)ethyl]acetamide (DIPPA) HCl (10 mg/kg Tocris) dissolved in 10% DMSO.

DOR agonist: (+)-4-[(αR)-α-((2S,5R)-4-Allyl-2,5-dimethyl-1-piperazinyl)-3-methoxybenzyl]-N,N-diethylbenzamide (SNC 80) (20mg/kg; Tocris) dissolved in 1mM HCl in saline.

DOR antagonist: naltrindole HCl (10 mg/kg; Tocris) dissolved in 10% DMSO.

All compounds were sonicated to ensure full dissolution and were administered sc. Dosages were adjusted for salt content where appropriate.

Vehicle: 10% DMSO in saline was chosen as the most appropriate vehicle solution.

Fluoxetine: The SSRI, fluoxetine, was included as a positive control for assay sensitivity. Fluoxetine (Sigma) was administered orally at 10 mg/kg suspended in maple syrup, daily for 4 weeks prior to testing and throughout the remaining duration of the study. Opioid receptor modulators and selected doses are summarized in table 1.

**Table 1:** FST scores

|  | **Immobility** | **Swimming** | **Climbing** |
| --- | --- | --- | --- |
| **IFN-α depression model** |  |  |  |
| Saline Control | 19.88 ± 1.5 | 19.5 ± 0.9 | 20.63 ± 2.1 |
| IFN-α + Vehicle | 36.88 ± 2.1*** | 12.63 ± 2.4* | 10.50 ± 1.9** |
| IFN-α + Fluoxetine (10mg/kg) | 28.50 ± 2.2<sup>#</sup> | 16.88 ± 1.1 | 14.63 ± 2.3 |
| ***Mu*** |  |  |  |
| IFN-α + Cyprodime (10mg/kg) | 34.75 ± 1.3 | 10 ± 1.7 | 15.25 ± 1.4 |
| IFN-α + RDC 2944 (0.1mg/kg) | 38.25 ± 2.9 | 12.0 ± 0.9 | 9.7 ± 2.4 |
| IFN-α + Morphine (5mg/kg) | 35.25 ± 1.0 | 20.25 ± 1.3 | 4.5 ± 0.5 |
| ***Kappa*** |  |  |  |
| IFN-α + DIPPA (10mg/kg) | 26.63 ± 1.2<sup>##</sup> | 15.7 ± 2.1 | 12.13 ± 2 |
| IFN-α + U50 488 (10mg/kg) | 38.13 ± 1.7 | 11.13 ± 1.1 | 9.6 ± 1.1 |
| ***Delta*** |  |  |  |
| IFN-α + Naltrindole (10mg/kg) | 30.38 ± 1.2 | 16.25 ± 1.9 | 16.6 ± 2.1 |
| IFN-α + SNC 80 (20mg/kg) | 19 ± 1.8<sup>###</sup> | 14 ± 1.2 | 25.75 ± 2.8<sup>###</sup> |
| IFN-α + DIPPA & SNC 80 | 21.13 ± 2.0<sup>###</sup> | 18.38 ± 1.7 | 16.38 ± 1.6 |

All drugs were a single administration, sc, except for fluoxetine, which was administered daily (po) for 4 weeks)
\* indicates significant difference to saline control group \*p<0.05, \*\*p<0.01, \*\*\*p<0.0001

#### 2.2.3. Experimental Procedures

##### 2.2.3.1. Forced swim test (FST)

The FST was conducted approximately 1hr (training) and 24hr (test) after IFN-α / saline treatment in relevant groups. The FST was used as a measure of stress coping behavior. On the training day rats were placed into the glass cylinders (30cm diameter) filled to a depth of 20cm with water (25±1°C) for 15min. On the test day 24hr after training, a 5min test was conducted. Behavior was video recorded and later scored by an observer blind to treatment. Immobility, climbing and swimming behaviors were scored by the count method as per Slattery and Cryan [28]. Opioid modulators or vehicle were administered 1hr before the FST.

##### 2.2.3.2. Progressive Ratio Schedule

The progressive ratio (PR) schedule was conducted similar to previous publications [29, 30]. Briefly, the PR schedule requires the animal to press a retractable lever presented on a random basis on the left or right side of the operant panel to initiate the trial, which is followed by delivery of a sucrose pellet into the hopper. Trials are separated by 5s. With each trial the number of lever presses required to earn a sucrose reward increases randomly by a factor of 3 or 5. Overall performance on the task declines in a linear manner as the PR increases. This is used as a measure of motivation, indicating how hard the animal is willing to work for a single sucrose pellet reward. Animals are maintained under a controlled food allowance of 50g lab chow per kg per rat, at the end of each day, for the duration of the experiment.

Apparatus: Experiments were performed in a Plexiglas behavioral testing chamber (Med Associates; 30.5 cm L x 24.1 cm W x 21.0 cm H), with two retractable levers mounted on one wall 5.5 cm above the chamber floor (distance between levers is 11 cm), a hopper (5 cm x 5 cm) mounted between the levers 1.5 cm above the floor on the original wall where 45 mg dustless sucrose pellets were delivered (TestDiet^TM^, 5TUT formula).

*Training Protocol:*

*Habituation*: Rats received 3 days of exposure to the chamber with levers retracted. During these 30min sessions, rats learned to consume sucrose pellets delivered every 20 ± 15s to the hopper.

*Continuous reinforcement (CRF) schedule*: Following habituation, the CRF program is implemented where, at random, the right or left lever is extended into the chamber and rats are trained to press the lever to receive a sucrose reinforcer. In each 30min session, sucrose pellets were delivered on a CRF schedule, where each lever press was followed by the delivery of a sucrose pellet. Additionally, a sucrose pellet was delivered every 120s, non-contingent on lever press behavior. Once rats made a minimum of 100 lever presses in a single session (approximately 3 sessions), they were moved to a fixed reinforcement schedule.

*Fixed reinforcement (FR) schedule*: Sessions were extended to 60min duration and reinforcement was delivered on a fixed-ratio 3 (FR3) schedule of reinforcement, for which reinforcement was delivered after every three lever presses. Non-contingent reward was discontinued during this and subsequent sessions. Once rats had made 100 lever presses in each of two sessions of FR3, rats were moved to progressive ratio training (approximately 4 sessions).

*PR training*: Rats underwent 30min sessions of lever press training on a progressive ratio schedule of reinforcement. On a PR schedule, the number of lever presses required for delivery of reinforcement increased by 3 or 5, at random, with each additional trial. Training continued for 10 days.

On the 11^th^ day, 30min before entering the operant chambers, animals were administered an opioid receptor modulator or a final fluoxetine treatment. Animals were then tested in the progressive ratio schedule at 30min and 24hr post drug and repeatedly tested for 6 days post drug. IFN-α / saline was administered approximately 24hr before the opioid receptor modulator (1hr before tests 2, 5 and 7) to reduce the risk of drug-drug interaction.

### 2.3. Neurogenesis, Sample Preparation and Tissue Collection

BrdU was administered once daily for four consecutive days on PR schedule test days 4, 5, 6 and 7 (days 52 to 55 in the overall timeline, Figure 2) at 100 mg/kg ip (Sigma), at approximately 17:00 hr. Sixteen hours later, animals were sacrificed. All rats were anesthetized by isoflurane inhalation and decapitated, and hippocampal samples were extracted for flow cytometric analysis of neurogenesis. The analysis of BrdU-labelled cells was undertaken as per previously published method using FITC BrdU Flowkit (BD Pharmingen™ #559619) [15, 31].

### 2.4. Statistical analysis

Statistical analyses were performed using Graphpad 6 and figures were constructed using CoralDraw X6. FST and BrdU+ cells data were analyzed as follows: comparisons between saline-controls and vehicle-IFN-α animals was by students t-test, comparisons between all IFN-α treated groups was by one-way analyses of variance (ANOVA) followed by Bonferroni *post hoc*, when appropriate. Analysis of data from the PR schedule consisted of within group analysis by one-way repeated measures (RM) ANOVA followed, when appropriate by Bonferroni *post hoc* analysis. Between group analysis was by two-way RM ANOVA followed, when appropriate by Bonferroni *post hoc* analysis. Cohen’s d was calculated as a measure of standardized effect size [32]. Small, medium, and large effect sizes in animal studies are equivalent to d values of 0.5, 1.0, and 1.5, respectively. Effect sizes were included to characterize the meaningfulness of a group difference in this behavioral study. An effect size, in the context of this study, also can be understood as the average percentile of the average drug-plus IFN-α-treated rat relative to the average vehicle-treated rat.

## 3. Results

### 3.1. Effects of single dose opioid receptor modulator or chronic fluoxetine on IFN-α-induced increase of immobility behavior in the FST

IFN-α-treated rats had significantly higher (Table 1) immobility scores than saline controls (t test: t = 6.4, p < 0.0001), and significantly lower swimming and climbing scores than saline controls (t tests: t = 2.67, p = 0.018 and t = 3.49, p = 0.003, respectively). Immobility data are represented as a percentage of saline control, in order of effect size (Figure 3). Acute treatment with SNC 80, DIPPA or a combination of the two have similar anti-immobility effects as chronic fluoxetine treatment. Cohen’s d relative to the IFN-α group was 1.35, 2.16, 2.68 and 3.16, indicating a medium to large effect size for fluoxetine, DIPPA, SNC 80 plus DIPPA and SNC 80 alone (Figure 3).

**Figure 3.**
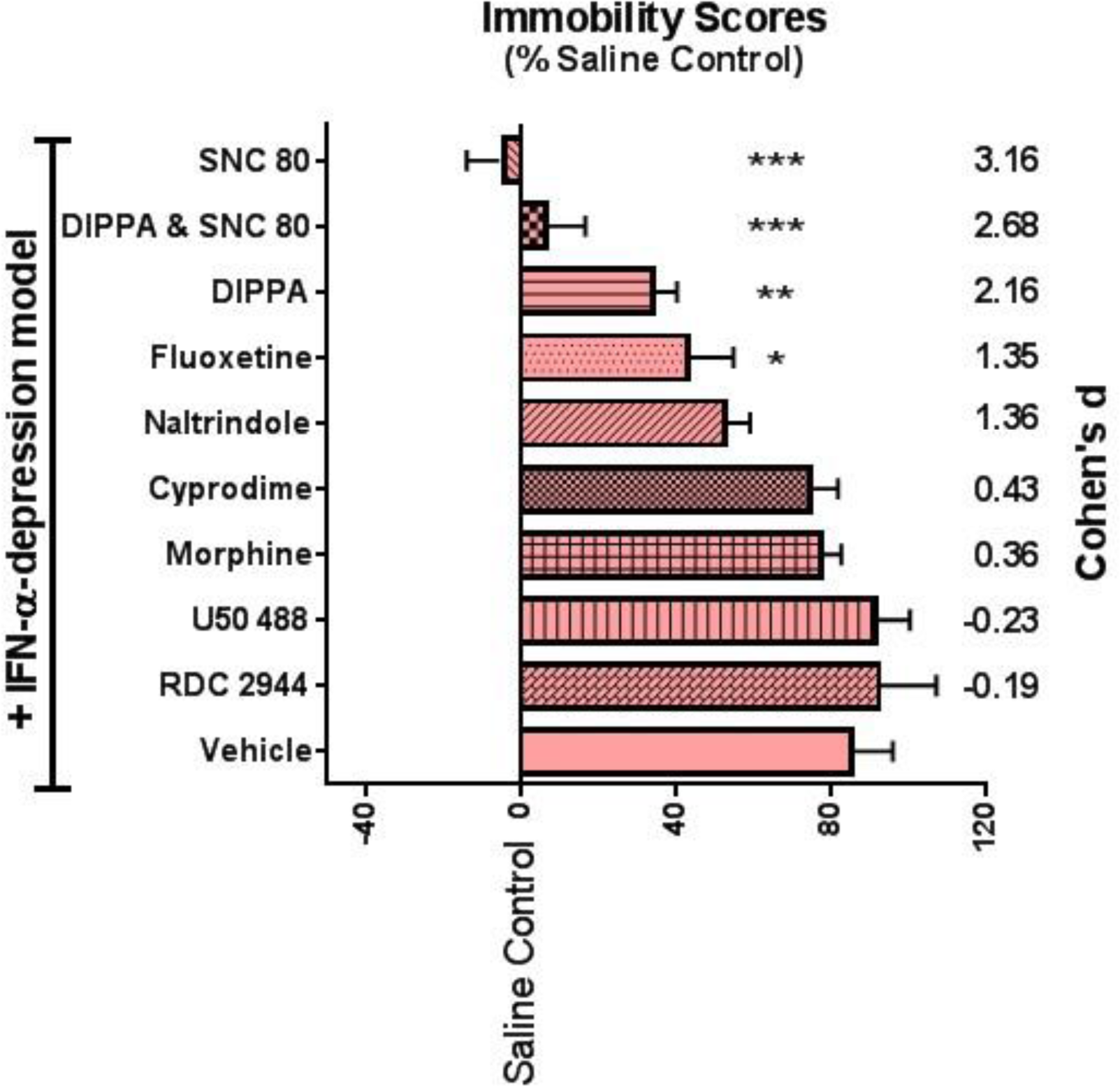
Acute treatment with SNC 80 or DIPPA or a combination of the two are comparable to chronic fluoxetine treatment on immobility behavior in the FST in IFN-α depression model. Cohen’s d values indicate the effect size compared to vehicle treated IFN-α rats. Histogram represent data expressed as a percentage of the saline control group (mean ± SEM), n=8 per group, asterisk represents significant difference to vehicle IFN-α group. One-way ANOVA with Bonferroni *post hoc* analysis ***p<0.0001, **p<0.01, *p<0.05.

Comparison of immobility scores between IFN-α groups treated with vehicle, chronic fluoxetine or acute opioid receptor modulators indicated a significant difference between groups (all data in Table 1: ANOVA: F(9, 70) = 14.56, p < 0.0001). Bonferroni *post hoc* analysis revealed chronic fluoxetine treatment, or single administration of the KOR antagonist, DIPPA, or DOR agonist, SNC 80, or the combination of DIPPA and SNC 80 significantly decreased immobility scores compared to vehicle-treated IFN-α rats (p < 0.05, p < 0.01, p < 0.001 and p < 0.001, respectively).

Comparison of swimming scores between IFN-α groups treated with vehicle, chronic fluoxetine or acute opioid receptor modulators indicated a significant difference between groups (Table 1: ANOVA: F(9, 70) = 4.261, p = 0.0002). Bonferroni *post hoc* analysis revealed IFN-α rats treated with morphine had significantly higher swimming scores compared to vehicle-treated IFN-α rats (p < 0.05).

Comparison of climbing scores between IFN-α groups treated with vehicle, chronic fluoxetine or acute opioid receptor modulators indicated a significant difference between groups (Table 1: ANOVA: F(9, 70) = 9.328, p < 0.0001). Bonferroni *post hoc* analysis revealed SNC 80 increased climbing score in the FST in IFN-α treated rats (p < 0.001) compared to vehicle-treated IFN-α rats.

### 3.2. Effects of single-dose opioid receptor modulators or chronic fluoxetine treatment on IFN-α-induced increase of break point in the PR task

IFN-α rats exhibited increased lever press behavior “breakpoint” (maximum number of lever presses per sucrose pellet reinforcement) in the PR task, compared to saline controls (Figure 4A and Table 2: t tests: p < 0.05 for each group).

**Figure 4.**
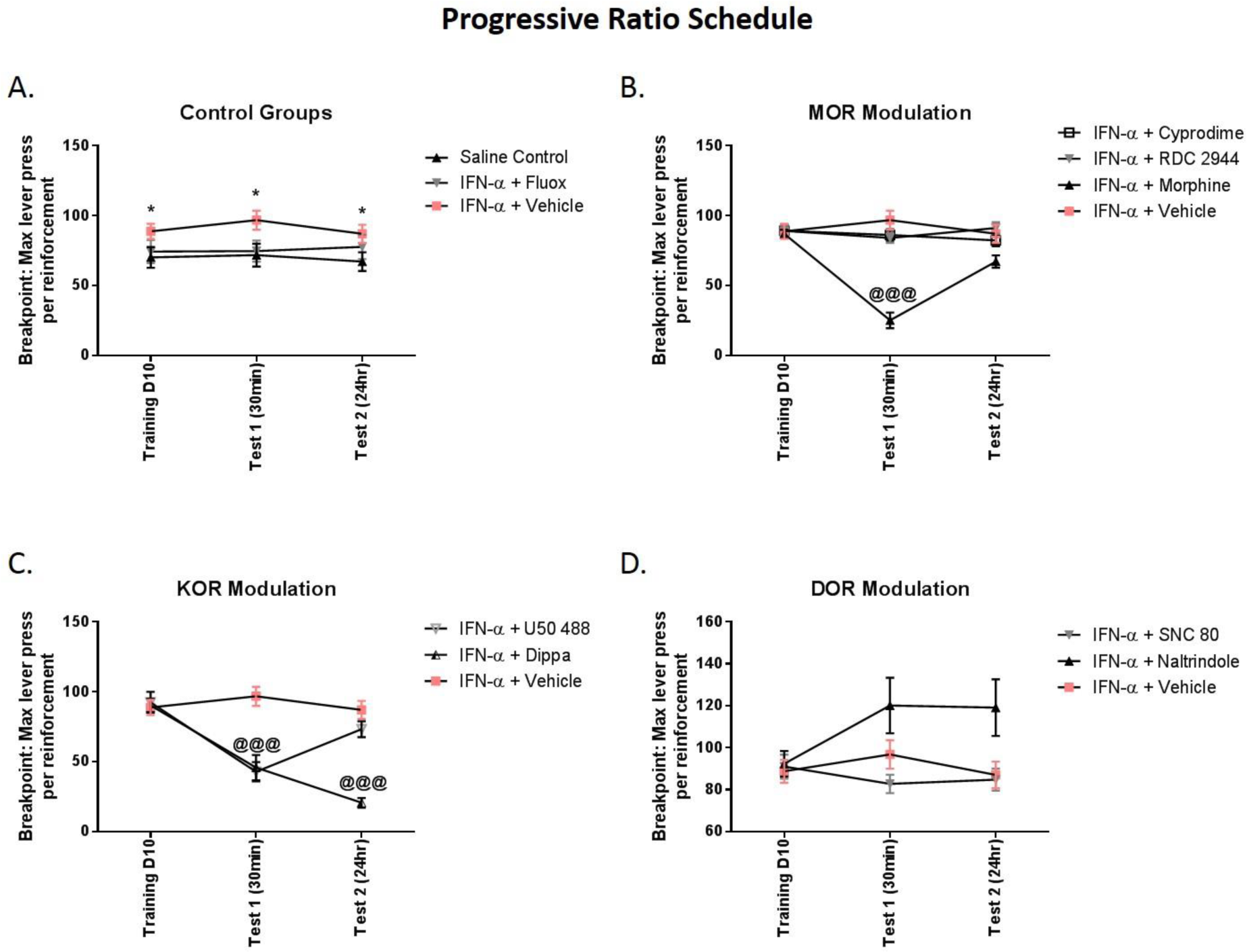
Acute treatment of IFN-α treated animals with morphine, DIPPA or U50 488 significantly decreased the breakpoint in the PR schedule compared to saline controls. A. IFN-α treatment induced an increase in the breakpoint point compared to saline controls *p<0.05, chronic fluoxetine treatment did not alter this. B. Acute treatment of IFN-α rats with morphine significantly reduced the breakpoint compared to vehicle treated IFN-α rats ^@@@^p<0.0001. C. Acute treatment of IFN-α rats with DIPPA or U50 488 significantly reduced the breakpoint compared to vehicle treated IFN-α rats ^@@@^p<0.0001. D. Acute treatment of IFN-α rats with naltrindole appears to increase the breakpoint in the PR schedule. Data are expressed as mean ± SEM, n=8 per group. Two-way RM ANOVA with Bonferroni *post hoc* analysis ^@@@^p<0.0001, ^@^p<0.05.

**Table 2:**
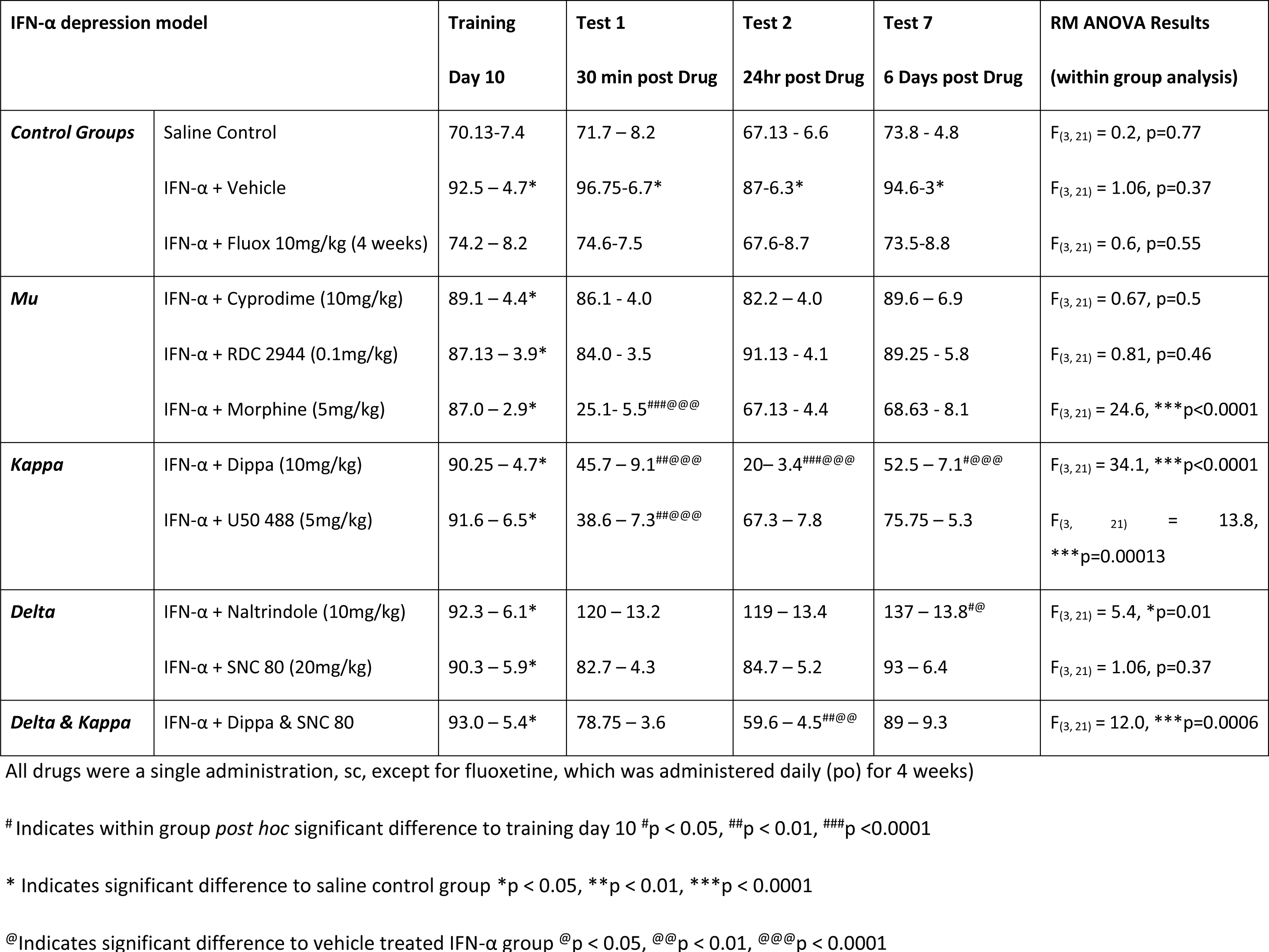
Progressive ratio schedule analysis

Treatment of IFN-α rats with fluoxetine for 4 weeks did not alter the performance in the PR when analyzed across test days (Figure 4A and Table 2: two-way RM ANOVA: treatment effect: F(1, 14) = 3.4, p = 0.08).

Treatment of IFN-α rats with the MOR antagonist, cyprodime (10mg/kg), or the low intrinsic activity MOR agonist, RDC 2944 (0.1mg/kg), did not alter performance in the PR task across test days (Figure 4B and Table 2). By contrast, a single dose of the MOR agonist, morphine (5mg/kg), significantly reduced the breakpoint in the PR task compared to vehicle treated-IFN-α rats (ANOVA, data in Table 2). Bonferroni *post hoc* analysis revealed the breakpoint was significantly reduced when tested 30min following morphine treatment, compared to training day 10 (p < 0.0001). This effect was absent at subsequent test times.

Treatment of IFN-α rats with a single dose of the KOR antagonist, DIPPA (10mg/kg), significantly reduced the breakpoint in the PR task (Figure 4C and ANOVA data in Table 2). Bonferroni *post hoc* analysis revealed the breakpoint was significantly reduced 30min and 24h post drug administration (p < 0.01, p < 0.0001, respectively), but not at post-treatment day 6 (p < 0.5). The KOR agonist, U50 488 (5mg/kg), also significantly changed the breakpoint in the PR task (data in Table 2). *Post hoc* analysis revealed the breakpoint was significantly reduced 30min following U50 488 treatment, compared to training day 10 (p < 0.01), but not at 24h and 6 days post dosing.

A single treatment of IFN-α rats with the DOR agonist, SNC 80, did not alter performance in the PR task (Figure 4D and ANOVA data in Table 2). Treatment of IFN-α rats with the DOR antagonist naltrindole (10mg/kg) significantly increased the breakpoint in the PR task (data in Table 2). *Post hoc* analysis revealed a significant increase in the breakpoint on day 6 post naltrindole administration, compared to day 10 training prior to drug treatment (p < 0.05).

Treatment of IFN-α rats with a combination of DIPPA and SNC 80 recovered the IFN-α-induced increase in the breakpoint in the PR task (ANOVA data in Table 2). Bonferroni *post hoc* analysis revealed a significant decrease in the breakpoint 24h post dosing (p < 0.01).

### 3.3. Effects of a single dose of opioid receptor modulators on IFN-α-induced decrease of hippocampal neurogenesis

Twenty four hr after completion of the last test in the PR schedule, animals were sacrificed, hippocampal tissue was extracted and processed for detection of BrdU-labelled cells as an indication of neurogenesis. IFN-α treatment decreased the number of hippocampal BrdU-labelled cells, compared to saline controls. This decrease in neurogenesis was reversed by a single-dose of SNC 80 (Figure 5a: ANOVA: F(11, 78) = 2.817, p = 0.0038). Bonferroni *post hoc* analysis revealed BrdU-labelled cells were significantly decreased in vehicle-treated IFN-α rats, compared to saline controls (p < 0.01), and this was recovered by treatment with SNC 80 alone (p < 0.01) or a combination of DIPPA and SNC 80 treatment (p < 0.05). Chronic fluoxetine treatment failed to recover IFN-α-induced decrease in BrdU-labelled cells in the group analysis.

**Figure 5.**
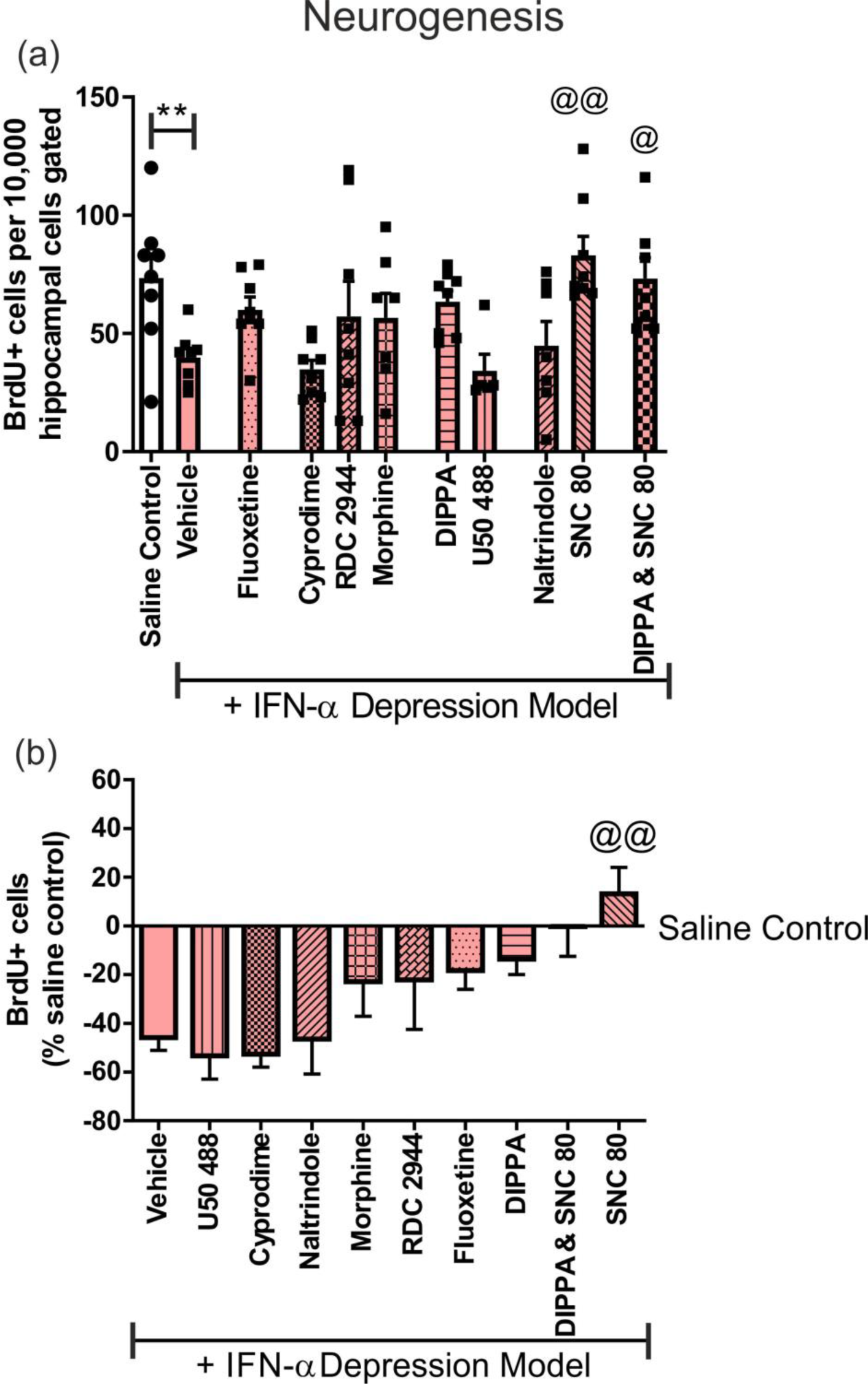
Acute treatment with SNC 80 or in combination with DIPPA recovered IFN-α-induced decrease in hippocampal BrdU-labelled cells. Histogram represents quantification of BrdU+ cells per 10,000 cells gated. (**a**) Acute treatment with the DOR agonist, SNC 80 alone, or in combination with the KOR antagonist, DIPPA, recovered IFN-α-induced reduction in hippocampal BrdU-labelled cells and chronic treatment with fluoxetine had no effect. (**b**) Histogram represent data expressed as a percentage of the saline control group. Acute treatment with the DOR agonist SNC 80 recovered IFN-α-induced reduction in hippocampal BrdU-labelled cells. Data are expressed as mean ± SEM, n=8 per group, * represents significant difference to saline control, @ represents significant difference to vehicle treated IFN-α group. Analysis by one-way ANOVA followed by Bonferroni *post hoc* tests **p<0.01, ^@^p<0.05, ^@@^p<0.01.

The data are also expressed as percent of saline control group (Figure 5b). This analysis revealed that IFN-α treatment reduced BrdU+ cells by approximately 40%, compared to saline controls. Chronic fluoxetine treatment recovers this effect by about 20%; only treatment with SNC 80 alone recovered the IFN-α-induced decrease in BrdU+ cells to surpass the saline control group, with an effect size of approximately 50%.

## 4. Discussion

The present study demonstrates single-dose acute treatment with the DOR agonist, SNC 80 or the KOR antagonist, DIPPA, or a combination treatment of the two, had anti-immobility effect in the FST comparable to chronic fluoxetine treatment in IFN-α-induced depression model. In the PR schedule, IFN-α-treated rats demonstrated an increased breakpoint compared to saline controls, which was reduced by acute treatment with the MOR agonist, morphine, KOR agonist U50 488 or KOR antagonist, DIPPA. Chronic fluoxetine treatment had no effect in the PR task. Acute treatment with the DOR antagonist naltrindole increased the breakpoint, compared to vehicle-treated IFN-α rats. SNC 80 alone or a combination of SNC 80 and DIPPA recovered the IFN-α induced decrease in hippocampal neurogenesis. Overall, these data implicate differential roles of MORs, KORs and DORs in the regulation of stress-coping and motivational behaviors and hippocampal neurogenesis in the back-translational rat IFN-α-induced depression model.

IFN-α-treated rats displayed increased immobility in the FST, corroborating our previous work [15–18], and chronic treatment with fluoxetine prevented this increase in immobility. We found no effect of MOR activation, since neither the full agonist, morphine, nor the low intrinsic activity agonist, RDC 2944, had an effect on IFN-α-induced increase of immobility. These findings differ from previous FST data showing anti-immobility effects of the potent MOR agonist and KOR antagonist, buprenorphine, in the Wistar-Kyoto rat [33], possibly indicating that different biological pathways underpin the increased immobility behavior in IFN-α and Wistar-Kyoto rats subjected to the forced swim stress. Another possibility is that KOR antagonism possibly in combination with MOR agonism mediates the effect of buprenorphine [33]. The lack of effect of morphine and RDC 2944 here may also be due to differences in the degree of MOR engagement produced by the doses applied, and the resulting activation of brain MORs. Further, both morphine and RDC 2944 were tested in the FST at 1hr following treatment, whereas buprenorphine was tested at 24hr following administration in the previous study [33], which may account for the different observations. We also report acute treatment with the KOR antagonist, DIPPA, had an anti-immobility effect in the IFN-α-treated rats, with a correspondingly large effect size. Previous work has shown the KOR antagonist, nor-BNI, produced robust anti-immobility effects in the Wistar-Kyoto rat [34]. Interestingly, KOR knockout mice showed no behavioral changes in the FST compared to wild-type mice [8]. This suggests that the antagonism of KORs engages signalling effects absent in the KO mice. We also found acute treatment with the DOR agonist, SNC 80, decreased immobility in IFN-α-treated rats, with a particularly large effect size. The peptide (DPDPE, JOM-13 and deltorphin II) and non-peptide (SNC 80 and (+)BW373U86) DOR agonists all show decreased immobility in the FST following acute or sub-chronic dosing [35–39]. A concern with DOR agonists is the potential pro-convulsant properties reported in rodents [40, 41]. We did not observe convulsions in any of the animals in our study. Treatment of IFN-α rats with the MOR antagonist, cyprodime, DOR antagonist naltrindole or KOR agonist U50 488 was without effect on immobility in the FST. In conclusion, only KOR antagonism or DOR agonism modulated stress-coping behavior in the FST in IFN-α treated rats. Since the doses of DIPPA and SNC 80 produced maximum or close to maximum responses in the FST, no additional effect could be found by combining them. A study of the KOR antagonist, LY2444296, and the DOR agonist, ADL5859, alone, and in combination yielded significant synergistic anti-immobility effects in the mouse FST [42]. However, this study used lower doses of the individual drugs than we employed which may allow for the synergistic effect observed.

We assessed motivational behavior using a PR schedule to index motivation to obtain sucrose reward. Interestingly, IFN-α treated rats had a higher breakpoint than controls, perhaps indicating a dysregulated reward pathway arising from IFN-α treatment. In a recent study, patients with hepatitis C viral (HCV) infection receiving IFN-α exhibited reduced activation of the ventral striatum in the win vs lose condition of a gambling task compared with patients with HCV awaiting treatment [43]. These data suggest IFN cytokine therapy can alter function of brain regions supporting motivation. In our study, chronic treatment with fluoxetine may have prevented IFN-α-induced dysregulation of the reward response in the PR task since both treatments were initiated at the start of the study and the PR response was comparable to vehicle control animals. It is unknown whether fluoxetine would have affected an already established IFN-α-induced motivational imbalance in our assay.

Acute treatment with morphine, DIPPA or U50 488 reduced the elevated breakpoint of IFN-α-treated rats in the PR schedule. The effects of morphine and U50 488 were only observed at the 30min test timepoint, at later time-points the effect disappeared. MORs have a complex relationship with hedonic and appetitive behaviors: for example, Zhang and colleagues demonstrated morphine withdrawal resulted in decreased motivation to obtain a natural reinforcement [44], while others showed morphine withdrawal increased operant responding for sucrose reward, but only for a 60% or greater sucrose solution [45]. These data suggest that morphine dosing, time of withdrawal and concentration of sucrose can easily alter results obtained. With respect to the effect of both morphine and U50 488 the doses at the 30min post dosing timepoint may have compromised motor functions, confounding data interpretation. Acute treatment with MOR or KOR agonists has previously been shown to increase locomotor activity [46, 47]. In a similar study the KOR agonist, salvinorin A (2mg/kg), also lowered breakpoint in a PR schedule in rats reinforced with a sucrose reward [48]. Although the authors did not measure locomotion, a previous study demonstrated salvinorin A at this dose did not alter locomotion [49]. Acute treatment with the KOR antagonist DIPPA decreased breakpoint in the PR schedule in IFN-α treated rats from 24h post drug administration, up to 6 days which is long after the compound has been eliminated from the body.

To our knowledge this is the first study which has examined the effects of a KOR antagonist on motivation to obtain a natural reinforcement. Interestingly, KORs can utilize different signalling cascades through GPCRs and β-arrestin, resulting in opposing stress-preventing or stress-elicited behavioral effects [50]. The prolonged duration of action of KOR antagonists has been attributed to c-Jun N-terminal kinase (JNK) 1 activation [51]. Controlled manipulation of KOR signalling may allow for development of therapeutics without the complications of dysphoria. The MOR antagonist and low intrinsic activity agonist, cyprodime and RDC 2944 respectively, had no effect on behavior in the PR schedule in the IFN-α depression model. The dose of RDC 2944 was 0.1mg/kg which corresponds to approximately 20% brain MOR occupancy, which is enough to induce pharmacological response in other MOR agonist related assays. This is much lower than the estimated MOR occupancy by the full agonist morphine at 5mg/kg and it cannot be excluded that a RDC 2944 dose targeting a higher level of MOR occupancy may have had effects similar to morphine.

The DOR agonist, SNC 80, also had no effect on breakpoint in IFN-α rats. There is little evidence in the literature to support a role for DORs in reward and motivational behaviors. Interestingly the DOR antagonist, naltrindole, increased the breakpoint further in the IFN-α rats 6 days post administration indicating that DOR modulation may be involved in the PR response in some way. It is believed that DORs contribute to contextual learning. Deletion of the DOR gene in mice impairs conditioned place learning but preserves morphine reinforcement, suggesting DORs do not contribute to opioid reward [52] but the context of learning reinforcement. The role of DORs in reward processing is not as clear as MORs and KORs. In conclusion, the involvement of opioid receptors in the regulation of IFN-α-induced dysregulation of motivational behavior is complex and may involve MOR, KOR as well as DOR. Studies involving isobolographic analysis may be warranted to understand the complex interaction between various opioid receptor mechanisms.

Neurogenesis is the process by which new neurons are generated in the hippocampus and is believed to facilitate plasticity [52]. Neurogenesis is increased by chronic treatment with SSRIs, and has been suggested as a marker of antidepressant activity in preclinical rodent models [53, 54]. We found IFN-α treatment suppressed cell proliferation in the hippocampal region [15, 55], and treatment with SNC 80 or a combination of DIPPA and SNC 80 reversed IFN-α-induced reduction of neurogenesis. The latter effect is likely ascribable to SNC 80 alone.

A single dose of each of the compounds selected to test was applied here; future studies including a range of different doses of the active compounds would be beneficial. Acute and chronic dosing regimens of select compounds would be interesting to compare results in the PR and the FST.

In conclusion, MOR, KOR and DOR mediate different aspects of IFN-α-induced dysregulation of motivational and stress-coping behavior and hippocampal neurogenesis. KOR and DOR appeared to play the more prominent roles. Notably, all effects were observed after a single dose of opioid receptor modulators; studies involving repeated dosing are needed to further explore the opioid modulation of stress and motivational behavior.

## Acknowledgements

We would like to thank Jean Bidlack and Lin E. Silver at the University of Rochester for the *in vitro* analysis of RDC 2944. We would also wish to thank Alan L. Pehrson at Montclair State University and Perry Cao (Alkermes, Inc.) for their contribution generating the ex vivo autoradiography and LC/MS/MS analysis respectively.

---

**Figure S1.**
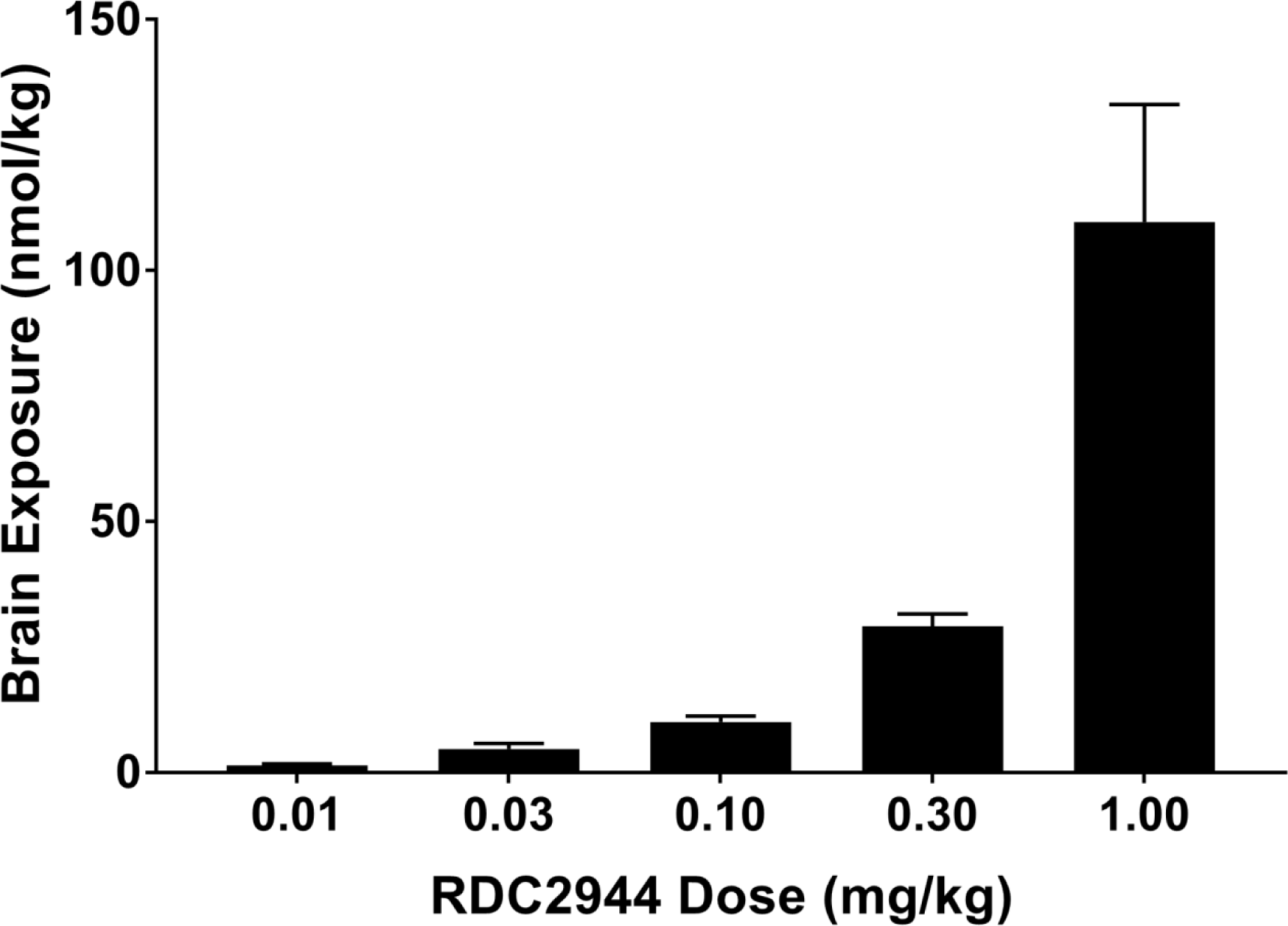
The relationship between RDC 2944 dose (1hr sc) and total brain exposure. Drugs were administered sc 1hr before sample collection. Dose was not corrected for salt. Data are expressed as mean + SEM.

**Figure S2.**
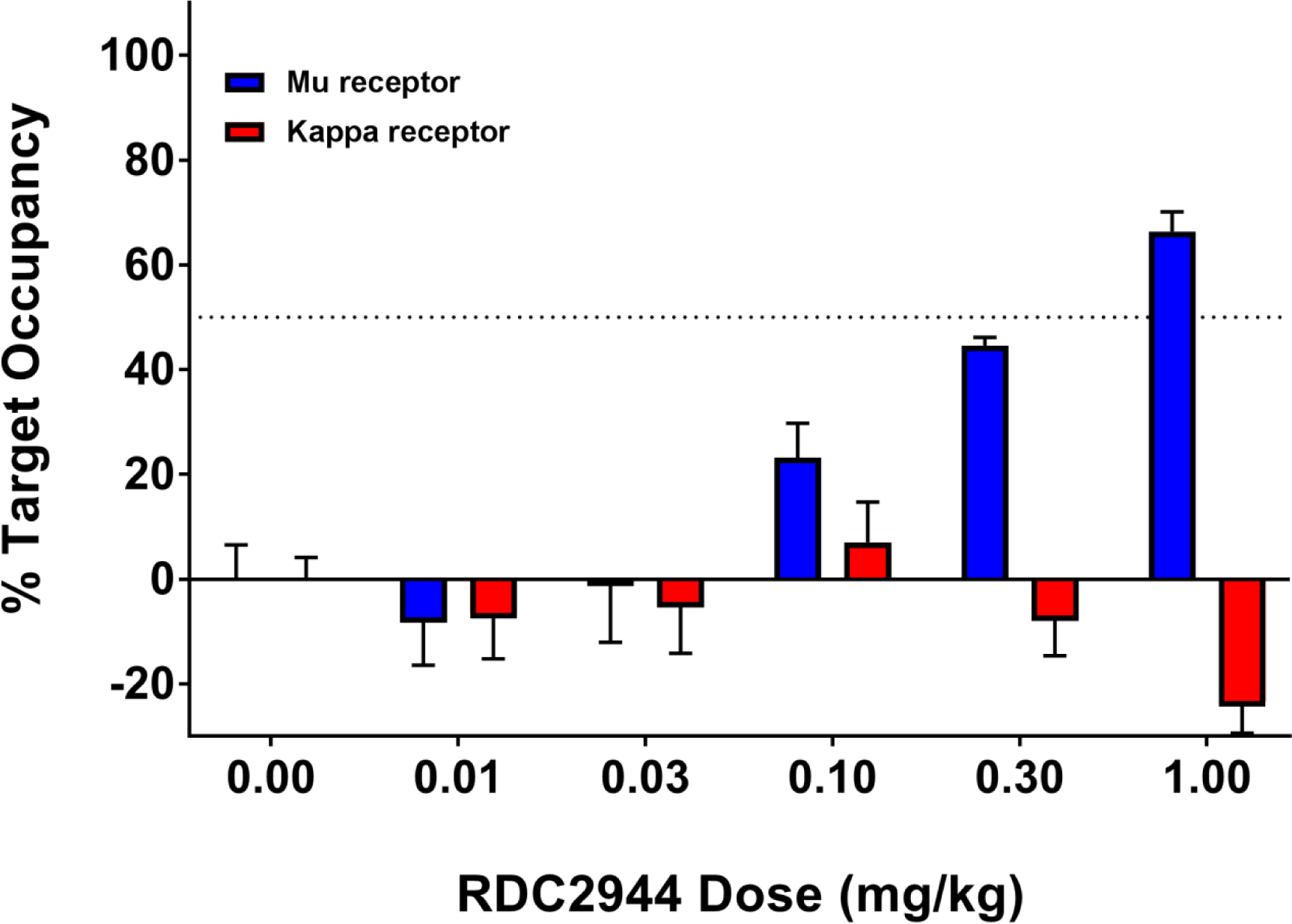
The relationship between RDC 2944 dose (1hr sc) and occupancy at µ and κ receptors. Drugs were administered sc 1hr before sample collection. Dose was not corrected for salt. Data are expressed as mean + SEM.

